# Contribution of the Wolffian duct mesenchyme to the formation of the female reproductive tract

**DOI:** 10.1101/2022.02.16.480738

**Authors:** Fei Zhao, Sara A Grimm, Shua Jia, Humphrey Hung-Chang Yao

## Abstract

The female reproductive tract develops from its embryonic precursor, the Müllerian duct.

In close proximity to the Müllerian duct lies the precursor for the male reproductive tract, the Wolffian duct, which is eliminated in the female embryo during sexual differentiation. We discovered that a component of the Wolffian duct, its mesenchyme, is not eliminated after sexual differentiation. Instead, the Wolffian duct mesenchyme underwent changes in transcriptome and chromatin accessibility from male tract to female tract identity, and became a unique mesenchymal population in the female reproductive tract with localization and transcriptome distinct from the mesenchyme derived from the Müllerian duct. Partial ablation of the Wolffian duct mesenchyme stunted the growth of the fetal female reproductive tract. These findings reveal a new fetal origin of mesenchymal tissues for female reproductive tract formation and reshape our understanding of sexual differentiation of reproductive tracts.

**Significance statement:** A female embryo initially possesses both the primitive female and male reproductive tracts, also known as the Müllerian and Wolffian duct. During sexual differentiation, the female eliminates the Wolffian duct and maintains the Müllerian duct that eventually differentiates into the female reproductive tract organs. However, in this paper, we show the female embryo retains mesenchymal cells surrounding the Wolffian duct for its reproductive tract formation. When incorporated into the female reproductive tract organs, the Wolffian duct mesenchyme shows unique anatomical localization and transcriptome and plays critical roles in female reproductive tract growth. This discovery provides new insights into female reproductive tract development and advances our understanding of sexual differentiation of reproductive tracts.

## Introduction

Sexually dimorphic differentiation of reproductive tracts in the mammalian embryo ensures the establishment of a sex-specific reproductive tract in adulthood (*1–3*). Prior to sexual differentiation, male and female embryos possess both primitive male and female reproductive tracts, known as the Wolffian and Müllerian ducts, respectively (*4*). During sexual differentiation, in the female embryo, the Wolffian duct regresses in the absence of androgens; whereas the Müllerian duct is maintained and eventually gives rise to the oviduct, uterus, cervix and upper vagina (*1*). In the male embryo, on the other hand, fetal testes produce two hormones to reverse the fates of the two primitive reproductive tracts: anti-Müllerian hormone (AMH) induces regression of the Müllerian duct while androgens maintain the Wolffian duct, which eventually differentiates into the epididymis, vas deference and seminal vesicle (*5*).

Prior to sexual differentiation (i.e. before embryonic day or E13.5 in mice), the Wolffian duct provides critical guidance for the formation of the Müllerian duct (*4, 6–9*). The Wolffian duct derives from the intermediate mesoderm and extends caudally till its distal part fuses with the urogenital sinus (*4, 8*). Shortly after the Wolffian duct formation, coelomic epithelium in the cranial part of the mesonephros is specified to become Müllerian duct precursor cells, which then invaginate caudally towards the Wolffian duct and form a canalized tube (*7*). Once in contact with the Wolffian duct, the Müllerian duct begins to elongate in the craniocaudal direction under the guidance of the pre-established Wolffian duct and eventually fuses at the urogenital sinus by E13.5 in mice (*6, 10*). The elongation of the Müllerian duct requires the activation of PI3K/AKT signaling pathway in rat Müllerian duct epithelia(*9*). It was thought that specification and invagination of specified cells in the Müllerian duct are independent of the Wolffian duct (*4*). However, a study in chicken embryos found that surgical removal of the cranial portion of the Wolffian duct before the onset of Müllerian duct formation abolished the expression of *Lhx1* (also known as *Lim1*) (*11*), the essential gene for the specification of coelomic epithelium to become Müllerian duct precursor cells (*12*). Therefore, the role of the Wolffian duct in specification and invagination during Müllerian duct formation may require further investigations. Nevertheless, substantial evidence has demonstrated that the subsequent step of Müllerian duct formation, its elongation, is dependent upon structural supports and regulatory signaling of the Wolffian duct. The tip cells of the elongating Müllerian duct contact closely with the basal membrane of the Wolffian duct, which serves as a physical guide for the Müllerian duct elongation (*13*). Physical interruption of the Wolffian duct in chicken embryos and genetic ablation of the caudal portion of the Wolffian duct in murine embryos both led to the arrested elongation of the Müllerian duct at the point of the interruption (*6, 14*). In addition, the Wolffian duct specifically expresses and secrets *Wnt9b*, a critical paracrine factor, without which the extension of the Müllerian duct is arrested at the cranial-most portion (*15*). Despite of the guiding roles of the Wolffian duct on Müllerian duct formation, the Wolffian duct epithelium itself does not contribute to or transform into the Mullerian duct. When the Wolffian duct epithelium was permanently labeled with a reporter *LacZ* that is induced by the Wolffian duct epithelium specific Cre (*Hox7b-Cre*), the entire Müllerian duct remained *LacZ* negative (*6*).

Despite its involvement in initial formation of the female reproductive tract before the onset of sexual differentiation, it is not known whether the Wolffian duct plays any contributing roles to the development of the female reproductive tract after sexual differentiation. When the Müllerian duct is formed under the guidance of the Wolffian duct, the two ducts are closely adjacent to each other and are surrounded by their respective mesenchymes, which are distinct in terms of microscopic appearance, gene expression and hormonal responsiveness. In the female mouse embryo on E13.5, the Müllerian duct mesenchyme lacks any recognized pattern in arrangement while mesenchymal cells surrounding the Wolffian duct appear elongated and radially disposed around the Wolffian duct (*16*). In terms of gene expression, the Wolffian duct mesenchyme but not the Müllerian duct mesenchyme expresses androgen receptor (*Ar*) (*17, 18*) and *Gli1*, a readout gene of the morphogen sonic hedgehog (SHH) that is specifically secreted from the Wolffian duct epithelium (*19–21*). On the other hand, the Müllerian duct mesenchyme expresses *Amhr2*, the specific receptor for AMH (*22*), whereas the Wolffian duct mesenchyme does not (*17, 18*). As a result, in the female embryo, the Müllerian duct mesenchyme can respond to anti-Müllerian hormone (*23*) while the Wolffian duct mesenchyme possesses the capability of responding to exogenous androgens (*24–26*).

During sexual differentiation, the epithelium of the Wolffian duct is eliminated in the female embryo. However, the fate of the remaining mesenchyme surrounding the Wolffian duct in the female embryo after sexual differentiation is unclear. Extensive studies have demonstrated the essential roles of the mesenchyme in the growth and differentiation of female reproductive tract organs after its early formation (*4, 5, 27, 28*). Given the juxtaposition of the Wolffian duct to the Müllerian duct, we set up to investigate whether the Wolffian duct mesenchyme is a source of functional mesenchymal tissues in the female reproductive tract .

## Results

### The Wolffian duct mesenchyme contributes to mesenchymal tissues in the female reproductive tract

Wolffian duct mesenchyme and Müllerian duct mesenchyme in the mesonephros can be distinguished by their specific expression of *Gli1* and *Amhr2*, respectively, during fetal development (*19, 22*). We confirmed this expression pattern by RNA-scope (**Fig. S1A**) and *Amhr2-Cre; Rosa-tdTomato; Gli1-lacZ* double reporter mouse line (**Fig. S1B**). To investigate what the Wolffian duct mesenchyme becomes in XX embryos, we developed the *Gli1-CreER;Rosa-tdTomato* tamoxifen-inducible lineage tracing model, where the Gli1-positive Wolffian duct mesenchymal cells (*Gli1^+^)* were permanently labeled with red fluorescent protein tdTomato only at the time of tamoxifen injection **(Fig. 1)**. Tamoxifen injection was performed on E13.5 and E14.5 before Wolffian duct regression in XX embryos. One day after the injection, a few mesenchymal cells surrounding the Wolffian duct in the cranial (future oviduct) and caudal (future uterus) portion of the mesonephros were positive for *tdTomato* (**Fig. 1A & F**). On E16.5 when the epithelium of the Wolffian duct disintegrated, the labeled *Gli1^+^* Wolffian duct mesenchyme remained present in the female reproductive tract **(Fig. 1B, 1C, 1G & 1H**). On postnatal day 0 (PND0 or birth), PND21 when the basic structures of the oviduct and uterus is established (*29*), and PND56 when the females are sexually mature for supporting pregnancy, the presence of *Gli1^+^* Wolffian duct mesenchyme persisted in the oviduct and uterus (**Fig. 1D, E, I & J**). These *Gli1^+^* Wolffian duct mesenchyme-derived cells expressed the markers for fibroblasts (Vimentin) (*30*) and smooth muscle cells (smooth muscle α-actin, αSMA) (*31*) (**Fig. S2**), indicating that they differentiated into typical cell types in the mesenchymal compartment of the female reproductive tract. These results demonstrate that the Wolffian duct mesenchyme in the XX embryo contributes to mesenchymal tissues in the female reproductive tract.

**Figure 1.**
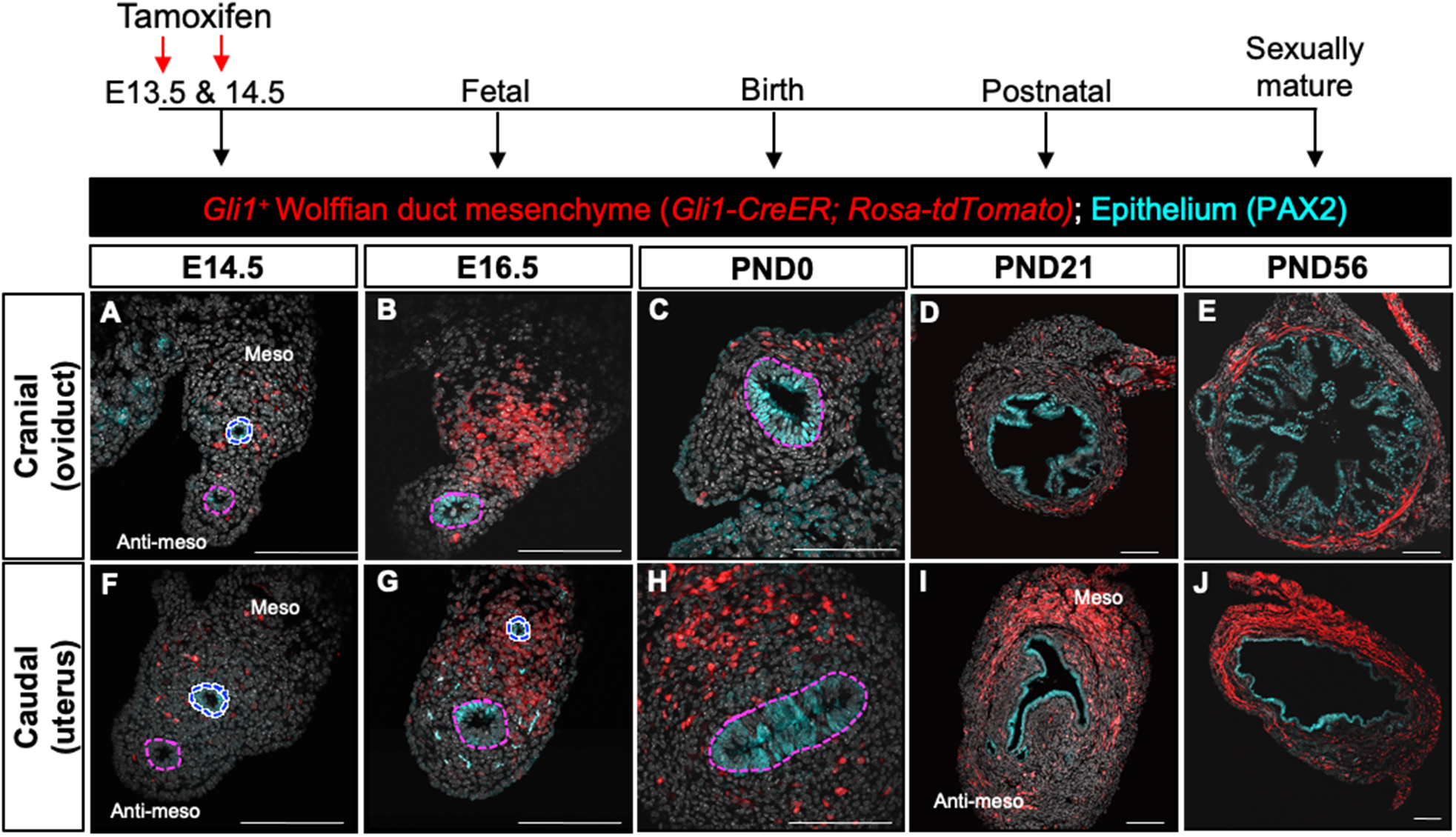
Lineage tracing of the *Gli1^+^* Wolffian duct mesenchyme in the female embryo. Tamoxifen was injected on E13.5 and E14.5 to permanently label *Gli1^+^* cells with tdTomato in *Gli1-CreER; Rosa-tdTomato* XX embryos. (**A-E**) cranial and (**F-J**) caudal sections of the female reproductive tract from tamoxifen treated females on E14.5 (**A & F**), E16.5 (**B & G**), PND0 (**C & H**), PND21 (**D & I**) and PND56 (**E & J**) to visualize *tdTomato*-labeled *Gli1^+^* cells (Red). All images are oriented with the mesometrial side of the tissues up and the anti-mesometrial side down. Epithelial cells are stained with PAX2 (Cyan). Pink and blue dashed lines circle Müllerian and Wolffian ducts, respectively. Scale bars in sections: 50 μm. N=3 in each time point examined.

### Transcriptomic and epigenetic changes underlying sexually dimorphic differentiation of the Wolffian duct mesenchyme

The Wolffian duct mesenchyme is masculinized to become components of the male reproductive tract in normal XY embryos or XX embryos exposed to exogenous androgens (*25, 32, 33*). Our discovery of the contribution of Wolffian duct mesenchyme to the female reproductive tract further indicates that the Wolffian duct mesenchyme is bipotential with sexually dimorphic differentiation capacity (**Fig. 2A**). We then set out to understand molecular changes and potential transcriptional regulation of the Wolffian duct mesenchyme when it follows either male or female fate. We performed RNA-seq and ATAC-seq to profile the transcriptome and chromatin accessibility landscape, respectively, on isolated *Gli1^+^*Wolffian duct mesenchymal cells from XX and XY mesonephroi on E14.5 and E16.5, the window of sexual differentiation of reproductive tracts. We plotted principal component analyses (PCA) to illustrate the degrees of differences among these four groups in terms of transcriptome and chromatin accessibility (**Fig. 2B & 2C**). From E14.5 to E16.5, the distances between XX and XY groups in both PCA plots were increased, suggesting that transcriptome and chromatin accessibility landscape of the Wolffian duct mesenchyme became more sexually divergent. These results were consistent the higher number of differential expressed genes (DEG) between XX and XY from E14.5 to E16.5 (**Fig. 2D & 2G**) and the increased number of differential ATAC-seq peaks (**Supplementary table 1**).

**Figure 2.**
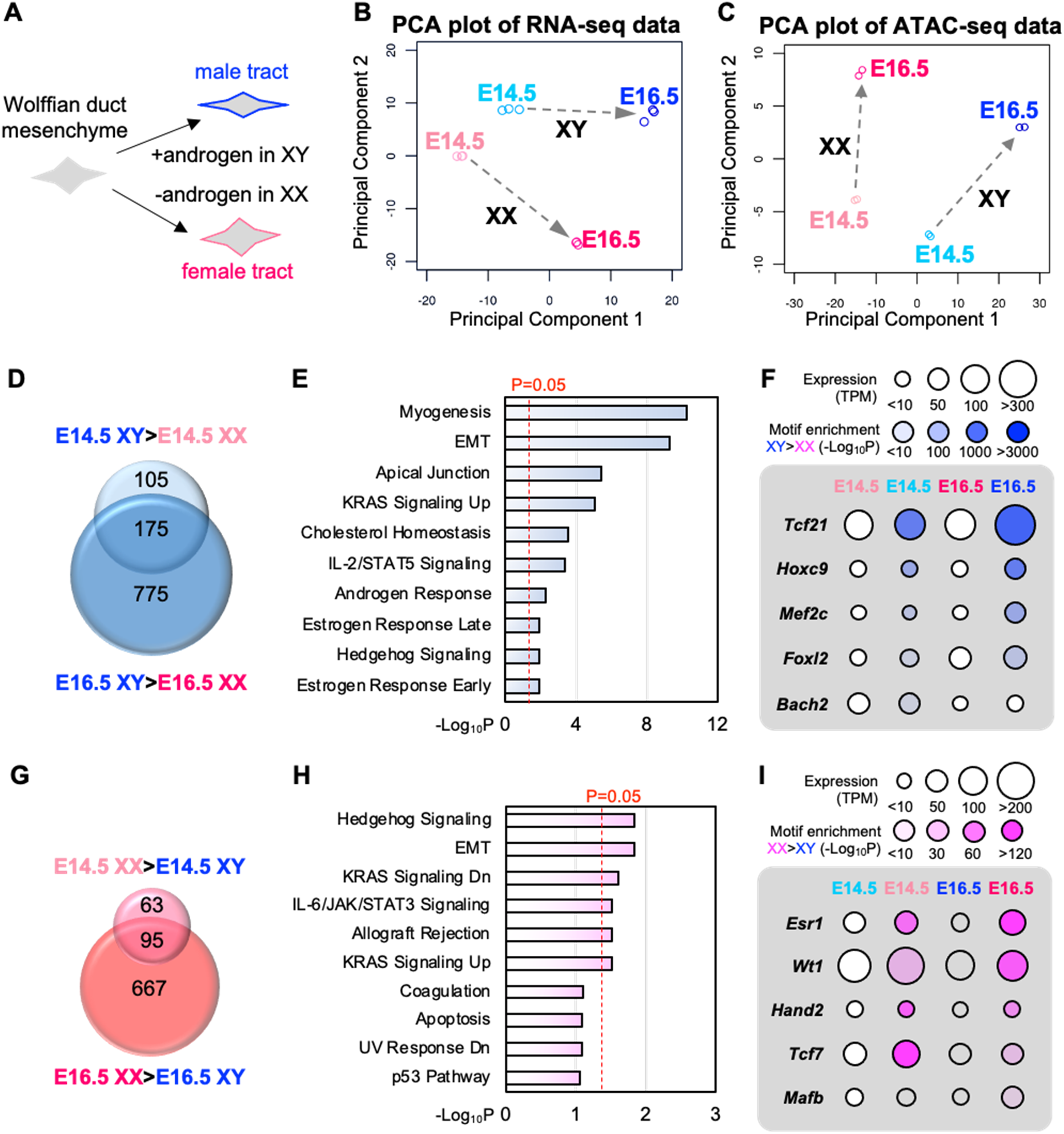
Profiling transcriptome and chromatin accessibility and deducing associated transcription factors in sexual differentiation of the *Gli1^+^* Wolffian duct mesenchyme. (**A**) The sexually dimorphic fate of the Wolffian duct mesenchyme in the presence (XY embryos) and absence (XX embryos) of androgens. (**B**) Principal component analysis (PCA) of the 500 most variant genes in the RNA-seq dataset including E14.5 XX (lighter red), E14.5 XY (lighter blue), E16.5 XX (bright red) and E16.5 XY (bright blue) *Gli1^+^* cells. N=3 in each group. (**C**) PCA of the 500 peak regions with the most variant signal in our ATAC-seq dataset. N=2 in each group. (**D & G**) Venn diagrams comparing significantly upregulated genes in XY (**D**) or XX (**G**) on E14.5 and E16.5. (**E & G**) Gene enrichment analyses of the upregulated genes in XY (**E**) (E14.5 XY>E14.5 XX or E16.5 XY>E16.5 XX) and in XX (**G**) (E14.5 XX > E14.5 XY or E16.5 XX > E16.5 XY). (**F & I**) Transcription factors (TF) which are in the list of upregulated genes in XY (**D**) and XX (**G**) and whose motifs are enriched in regions with increased chromatin accessibility in XY (**F**) and XX (**I**) in their comparisons on either E14.5 or E16.5.

To deduce biological events underlying male and female differentiation of the *Gli1^+^* Wolffian duct mesenchyme, we performed gene enrichment analysis of upregulated genes in XY and XX when they are compared (**Fig. 2D & 2G**). In these upregulated genes in XY *Gli1^+^* Wolffian duct mesenchymal cells (280 genes from E14.5 XY > E14.5 XX and 950 genes from E16.5 XY> E16.5 XX with 175 genes in common between the two stages), myogenesis, epithelial-mesenchymal transition and apical junction and KRAS signaling were significantly enriched with myogenesis or smooth muscle cell differentiation being the most significant enriched (**Fig. 2E**). This observation was consistent with the notion that Wolffian duct masculinization was associated with smooth muscle differentiation (*34*). On the other hand, when analyzing upregulated genes in XX *Gli1^+^* Wolffian duct mesenchymal cells (158 genes from E14.5 XX > E14.5 XY or 762 genes from E16.5 XX > E16.5 XY) (**Fig. 2G**), we found that the statistically significant pathways were Hedgehog signaling (a morphogen signaling), EMT (features extracellular matrix remodeling), KRAS signaling, IL-6/JAK/STAT3 signaling and allograft rejection signaling (**Fig. 2H**), which suggested remodeling in cellular morphology, extracellular matrix and immune responses during the female fate differentiation of the Wolffian duct mesenchyme.

At mRNA transcriptional level, actions of specific transcriptional factors (TFs) and accessible chromatin landscape for transcriptional factor bindings govern cellular state and differentiation (*35*). To identify transcriptional factors that potentially involve in male and female fate differentiation of the *Gli1+* Wolffian duct mesenchyme, we determined the upregulated TFs in XY or XX *Gli1^+^*Wolffian duct mesenchymal cells by overlapping the lists of upregulated genes in either of them with the mouse transcriptional factors database (*36*). We also examined TF binding motifs enriched in regions with increased chromatin accessibility in XY *Gli1^+^* Wolffian duct mesenchymal cells (E16.5 XY > E16.5 XX or E14.5 XY >14.5 XX) and XX (E16.5 XX > E16.5 XY or E14.5 XX > E14.5 XY) with the focus on those distal from transcription start sites, which contain a majority of differential peaks (85.7%, -95.5%, **Supplementary table 1**) and are expected to harbor TF-binding motifs critical for cellular differentiation (*37, 38*). The top 20 motifs enriched in regions of increased chromatin accessibility in XY and XX *Gli1^+^* Wolffian duct mesenchymal cells were shown in **Supplementary tables 2-5**. Finally, we overlapped the top motifs enriched at the chromatin regions with increased accessibility with the significantly upregulated TFs (TFs with the expression level TPM<10 were removed) in XY or XX *Gli1^+^* Wolffian duct mesenchymal cells.

For the male differentiation, this overlapping analysis yielded five transcriptional factors and their motifs, *Tcf21, Hoxc9, Mef2c, Foxl2,* and *Bach2* (**Fig. 2F**). Of note, *Ar* was not in in the list because *Ar* mRNA level in our RNA-seq datasets was not significantly different between XX and XY *Gli1+* Wolffian duct cells on either E14.5 or E16.5, which is consistent with previous findings (*17, 39*). However, androgen responsive elements (ARE) or AR-associated nuclear receptor motifs were the top motifs in regions with increased chromatin accessibility in XY *Gli1^+^* Wolffian duct mesenchymal cells. This observation demonstrates the benefits of including chromatin accessibility data for our analysis in identifying the predominant actions of AR. Among these five transcriptional factors, it has been shown that TCF21 modulates AR transcriptional regulation in vitro cancer cell lines (*40*); *Hoxc9* involves in regionalization of the Wolffian duct (*41*); and *Mef2c* (Myocyte-specific enhancer factor 2C) is a target of AR during androgen enhanced myogenesis (*42*). The roles of *Folx2 and Bach2* in Wolffian duct differentiation has not been reported in the literature. These results demonstrate that the male fate differentiation of the *Gli1^+^*Wolffian duct mesenchyme is associated with multiple TFs that are potential critical components in AR regulatory network.

We adopted the same strategy to infer potential TFs in the female fate differentiation of the *Gli1^+^* Wolffian duct mesenchyme (**Fig. 2I**). The upregulated genes in the XX *Gli1^+^* Wolffian duct mesenchymal cells included dozens of TFs, among which *Esr1*, *Wt1, Hand2*, *Etv4*, *Tcf7* and *Mafb* had their binding motifs enriched in regions with increased chromatin accessibility in XX (E16.5 XX > E16.5 XY or E14.5 XX > E14.5 XY) (Supplementary tables 4 **& 5**). The roles of *Wt1* and *Hand2* in fetal female reproductive tract development have not been reported. *Esr1* mediates estrogen actions and is critical for female reproductive tract function (*43*). However, the absence of *Esr1* does not affect female reproductive development (*44*). In the absence of either *Etv4* or *Tcf7*, female mice are fertile, suggesting that they play dispensable roles in the female reproductive tract development (*45, 46*). These observations demonstrate that the repurposing of the *Gli1^+^* Wolffian duct mesenchyme in XX embryos is associated actions of multiple transcriptional factors, which are more likely to be the outcome of the female fate differentiation.

### Similarities and differences in gene expression between *Gli1^+^* Wolffian duct mesenchyme and *Amhr2^+^* Müllerian tract mesenchyme in the neonatal uterus

Once becoming a part of the female reproductive tract, the *Gli1^+^*Wolffian duct mesenchyme-derived cells exhibited unique distribution along the tract: they were localized more in the uterus than the oviduct (**Fig. 1E** and **1F**). Additionally in the uterus, *Gli1^+^* Wolffian duct mesenchyme was clustered at the mesometrial side, where the uterine horn is connected to blood vessels and body cavity (**Fig. 1H & 1J**). Conversely, when we tracked the fate of the Müllerian duct-derived mesenchyme with the *Amhr2-Cre; Rosa-tdTomato* reporter model, we found that *Amhr2^+^* Müllerian tract mesenchyme contributed to the majority of mesenchymal cells in the oviduct and their localization in the uterus was exclusive to the anti-mesometrial side (**Fig. S3**). These results demonstrate that *Gli1^+^* Wolffian duct mesenchyme gives rise to a mesenchymal population distinct from *Amhr2^+^* Müllerian tract mesenchyme in the female reproductive tract.

To understand molecular similarities and differences between these two mesenchymal populations with their contrast localizations in the uterus (mesometrial vs anti-mesometrial side), we isolated these two cell types from the respective reporter models using FACS and compared their transcriptomic differences by RNA-seq at birth, when the Wolffian duct mesenchyme has been incorporated into the uterus. Both mesenchymal populations had high expression of typical mesenchymal genes, such as vimentin (*Vim*) and collagens (*Col1a1*, *Col1a2* and *Col3a1*). Nevertheless, their transcriptomes differed in 1,705 gene with 1049 genes expressed significantly higher in the *Gli1^+^* Wolffian duct mesenchyme and 656 genes higher in the *Amhr2^+^* Müllerian tract mesenchyme, respectively (**Fig. 3A**). To identify what signaling pathways have associations with these genes, we performing IPA upstream regulator analysis on the genes upregulated in each of the two mesenchymal cells. Interestingly, the analyses of higher expressed genes in these two populations showed enrichment of angiotensin (i.e. angiotensinogen and VEGF) and estrogen (i.e. beta-estradiol and diethylstilbestrol) signaling in common. Nevertheless, in the *Gli1^+^* Wolffian duct mesenchyme, the transforming growth factor beta one (TGFβ1) pathway was predicted to be one of the top signaling pathways that lie upstream of the 809 higher expressed genes. On the other hand, in the *Amhr2^+^* Müllerian tract mesenchyme, the WNT/beta-catenin or CTNNB1 pathways was enriched (**Fig. 3B**). Consistent with this analysis, a downstream target of TGFβ pathway *Akap12* (also known as SSeCKS) (*47*) was upregulated in the *Gli1^+^* Wolffian duct-derived mesenchymal cells and its protein was specifically localized to the Wolffian duct mesenchyme and the mesometrial side of the uterus on E16.5 and 18.5 (**Fig. 3C**). On the other hand, *Hand2,* a downstream target of WNT/beta-catenin signaling (*48*), was expressed higher in *Amhr2^+^*Müllerian tract mesenchyme and its protein was detected on the anti-mesometrial side of the E14.5-18.5 uterus (**Fig. 3D**). The enrichment of WNT/beta-catenin signaling pathway in *Amhr2^+^* Müllerian tract mesenchyme is consistent with the observation of higher *Wnt* signaling in uterine anti-mesometrial side (*49*). Taken together, our results unequivocally demonstrate the *Gli1^+^* Wolffian duct mesenchyme contributes a unique mesenchymal population different from the *Amhr2^+^* Müllerian duct mesenchymal cells in the female reproductive tract.

**Figure 3.**
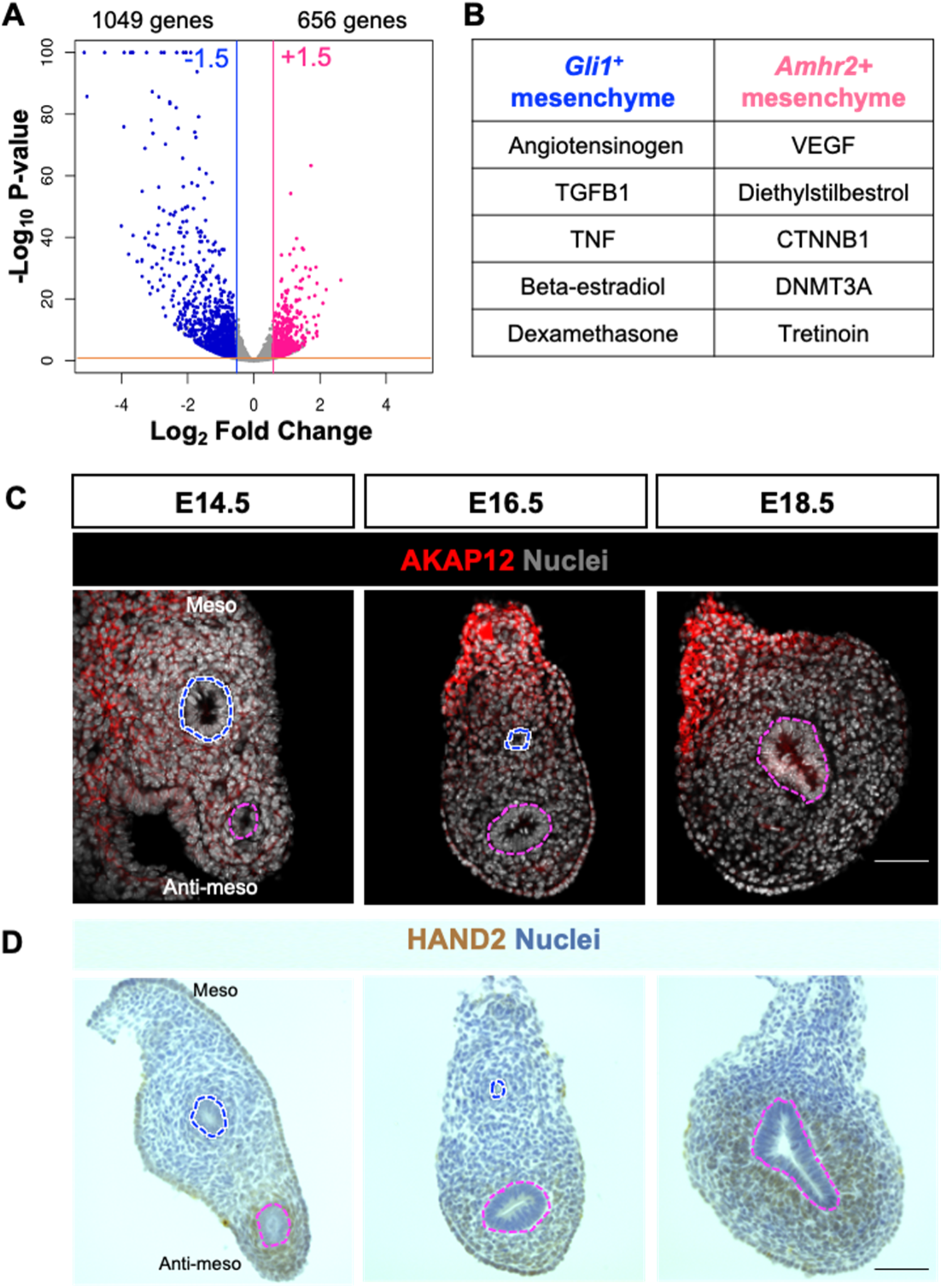
Gene expression similarities and differences between the *Amhr2^+^* Müllerian tract mesenchyme and *Gli1^+^* Wolffian duct mesenchyme in the neonatal uterus. (**A**) Volcano plot displaying differentially expressed genes between *Amhr2^+^* and *Gli1^+^* cells in neonatal uterus. N=3 in each group. A maximum value of 100 for -log10 (p-value) is assigned so that all genes are shown in the plot. Differentially expressed gene thresholds for significance and fold change are shown as solid lines. (**B**) The top five upstream regulators enriched in the higher expressed genes in *Amhr2^+^* and *Gli1^+^* cells, respectively. (**C**) Immunofluorescent staining of AKAP12 in the uterus on E14.5, E16.5 and E18.5; DAPI was used to stain nuclei (Grey). (**D**) Immunohistochemical staining of HAND2 in the uterus on E14.5, E16.5 and E18.5; Nuclei (blue) were stained with hematoxylin. All images in (**C & D**) are oriented with the mesometrial side of the tissues up and the anti-mesometrial side down. Pink and blue dashed lines circle Müllerian and Wolffian ducts, respectively. Scale bars in all sections: 50 μm. N=3 in each time point examined.

### Partial ablation of the Wolffian duct mesenchyme stunts the growth of female reproductive tracts ex vivo

Next, we investigated the functional significance of the *Gli1^+^* Wolffian duct mesenchyme in fetal female reproductive tract development by using the tamoxifen-inducible *Gli1-CreER; Rosa-tdTomato; Rosa-DTA* genetic cellular ablation model. In this model, the tamoxifen treatment not only induces *tdTomato* expression to label *Gli1^+^*cells, but also turns on the expression of diphtheria toxin or *DTA* that induces cellular apoptosis specifically in DTA-expressing cells (*50*). Because *Gli1* is expressed in mesenchymal compartments of other critical organs (*51*), *in vivo* cellular ablation of *Gli1^+^* cells led to multiple organ failures and embryonic lethality. To circumvent this problem, we performed the ablation experiment in organ culture, which is an established technique to study sexual differentiation of reproductive tracts ex vivo (*52*). Mesonephroi with *Gli1-CreER; Rosa-TdTomato* (control group) or *Gli1-CreER; Rosa-tdTomato; Rosa-DTA* (ablation group) from E14.5 XX embryos were cultured for two days with tamoxifen to induce *Cre* activities and tdTomato/DTA expression in the *Gli1^+^* Wolffian duct mesenchyme before Wolffian duct regression. Upon the tamoxifen treatment, *tdTomato* was still specifically expressed in the *Gli1^+^* Wolffian duct mesenchyme in both groups (**Fig. S4**). However, in the ablation group, we observed statistically significant increase in the number of apoptotic cells positive for the apoptotic marker, cleaved PARP1(*53*) in the mesenchyme (**Fig. S4)**. In addition, the total fluorescence signal of *tdTomato* in the mesonephroi after the culture was significantly reduced in the ablation group to 60% of that in the control group under the identical imaging setting **(Fig. 4A & C)**, indicating a partial ablation of the *Gli1^+^* Wolffian duct mesenchyme in the ablation group. Despite that the ablation was partial, we observed a noticeable phenotype that the lumen area of the Müllerian duct in the ablation group was significantly smaller than those in the control (**Fig. 4A & D**).

**Figure 4.**
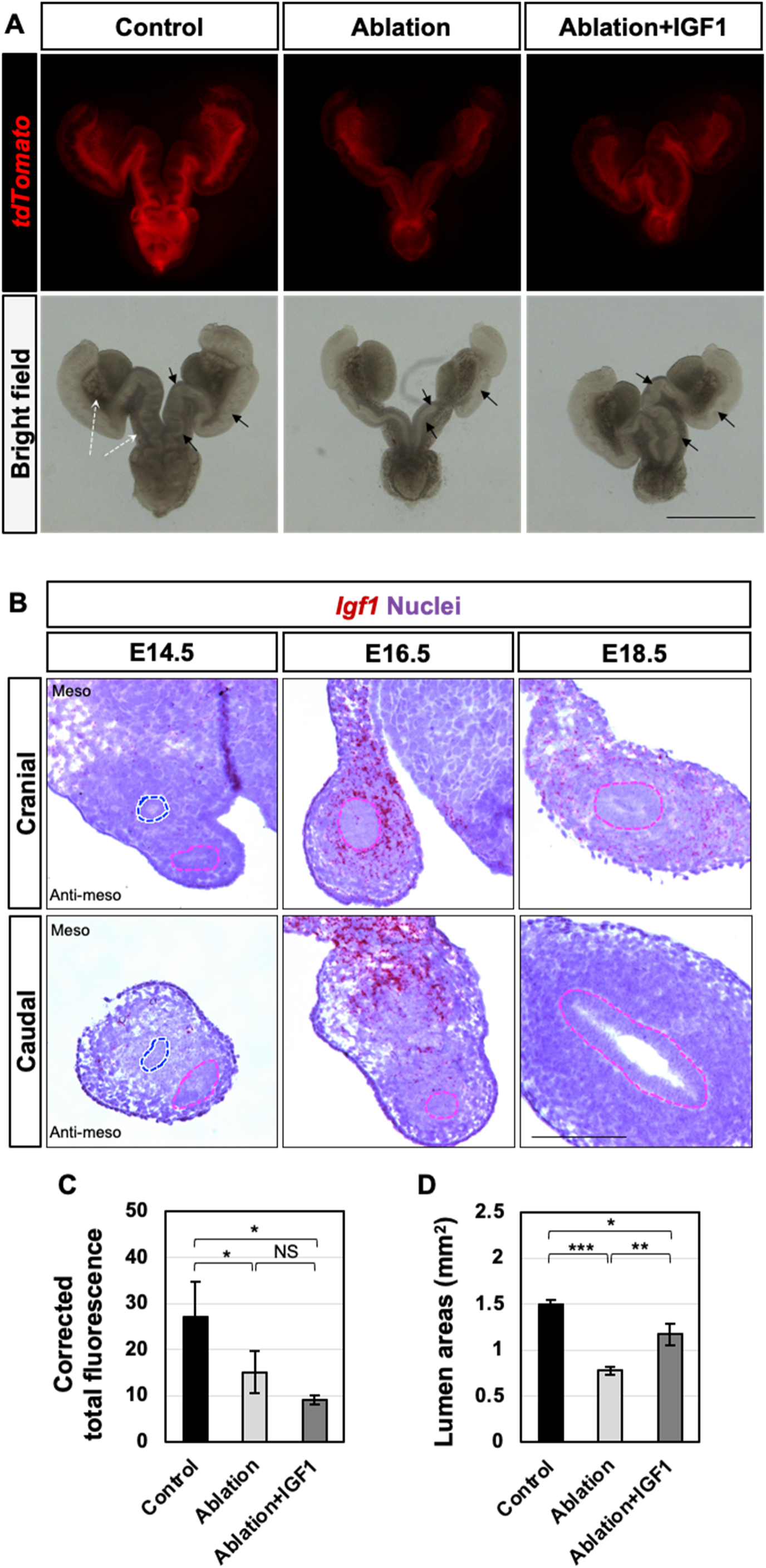
Partial ablation of the *Gli1^+^* Wolffian duct mesenchyme stunts the growth of female reproductive tracts ex vivo. (**A**) Whole mount *tdTomato* and bright-field images of the female reproductive tract with ovaries attached in the control (*Gli1-CreER+; Rosa-tdTomato+*), ablation (*Gli1-CreER+; Rosa-tdTomato+; Rosa-DTA)* and ablation+IGF1 (*Gli1-CreER+; Rosa-tdTomato+; Rosa-DTA* with IGF1 treatment) groups after 4-day culture. Scale bars: 1.4 mm. White and black arrows indicate the mesometrial side and Müllerian tract lumen, respectively. (**B**) Detection of *Igf1* in the cranial and caudal sections of the fetal female reproductive tract on E14.5, E16.5 and E18.5 by RNAscope. Pink and blue dashed lines circle Müllerian and Wolffian ducts, respectively. Scale bars: 100 μm. (**C**) Corrected total fluorescence of cultured female reproductive tracts in control, ablation, and ablation+IGF1 groups. NS: No significantly difference. *: P<0.05. (**D**) Lumen areas (mm^2^) of the fetal female reproductive tracts in the control, ablation, and ablation+IGF1 groups. N=4 for the control and ablation+IGF1 groups; N=5 for the ablation group. *: P<0.05; **: P<0.01; ***: P<0.001.

Next, we investigated how the partial loss of the *Gli1^+^* Wolffian duct mesenchyme led to the reduced lumen areas. It is well established that the paracrine growth factors from the mesenchyme regulates the growth and differentiation of the epithelium (*54*). Therefore, we searched for known growth factors (*55*) whose expression was significantly higher in the *Gli1^+^* Wolffian duct mesenchyme than in the *Amhr2^+^*Müllerian duct mesenchyme in our RNA-seq data (**Fig. 3A**). We then examined the expression pattern of the potential growth factors in fetal female reproductive tract and mouse knockout phenotypes in the Müllerian duct development through the online gene expression and mouse genome informatics databases (*56*). As a result, we uncovered that growth factors (*Angpt1, Angpt2, Bdnf, Fgf7*, *Fgf12, Igf1, Pdgfa,* and *Vegfd)* were enriched in the *Gli1^+^* Wolffian duct mesenchyme compared to the *Amhr2^+^* mesenchymal cells.

Among these factors, the mesenchyme-derived growth factor *Igf1* appeared to be a putative candidate with its receptor *Igfr1* expressed in the Müllerian duct epithelium (*57, 58*). Absence of *Igf1* or its receptor *Igf1r* led to hypoplastic reproductive tract organs in the XX embryo (*58, 59*), indicating the critical roles of *lgf1* signaling in the Müllerian duct growth. When we examined spatiotemporal expression of *Igf1* in the normal XX embryo, we found its specific and transient expression in the Wolffian duct mesenchyme at the mesometrial side of the caudal mesonephros (future uterus) on E16.5 although it was also expressed in the anti-mesometrial side of the cranial mesonephros (future oviduct) (**Fig. 4B**). These observations prompted us to investigate whether IGF1 was able to rescue the phenotype of the stunted growth of the Müllerian duct in the ablation group, where the ablation of *Gli1^+^* mesenchymal cells could lead to the decrease in mesenchymal IGF1 production. We supplemented exogenous IGF1 in the media of the ablation group and found that IGF1 treatment did not affect the ablation efficacy in the ablation groups (**Fig. 4A & 4C**).

However, the phenotype of the decreased lumen areas was partially rescued in the ablation group with IGF1 treatment (**Fig. 4A & 4D**). Taken together, these results indicate the *Gli1^+^* Wolffian duct mesenchyme is critical for the proper growth of the fetal female reproductive tract, probably through the paracrine growth factor IGF1.

## Discussion

We provide the molecular and genetic evidence for the dual origins of mesenchymal tissues in the female reproductive tract. We used *in vivo* genetic cell tracking approach to reveal that the *Gli1^+^* Wolffian duct mesenchyme continues to exist in the developing female reproductive tract, corroborating with another study with slightly different approach (*60*). We also revealed transcriptome and chromatin accessibility changes in sexual differentiation of the *Gli1^+^* Wolffian duct mesenchyme, transcriptomic similarities and differences between the *Gli1^+^* Wolffian duct mesenchyme and the *Amhr2^+^*Müllerian duct mesenchyme, and the functional significance of the *Gli1^+^* Wolffian duct mesenchyme in the fetal growth of the female reproductive tract (**Fig. 5**).

**Figure 5.**
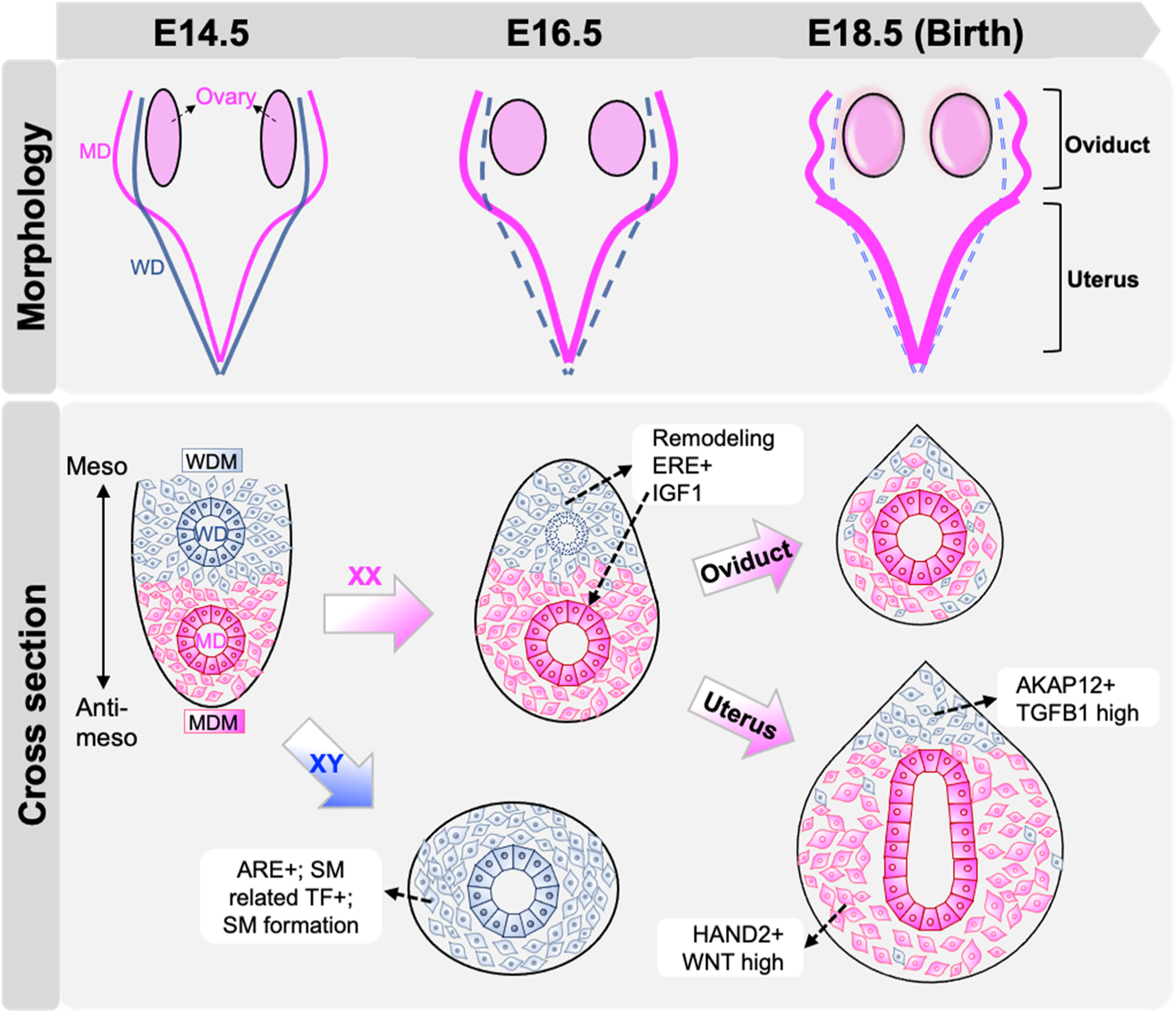
The model summarizing the major results on the fates and roles of the *Gli1^+^* Wolffian duct mesenchyme in reproductive tract development. At the onset of sexual differentiation of reproductive tracts (E14.5 in the mouse), the XX embryo possesses both the Wolffian duct (WD) and the Müllerian duct (MD), which are surrounded by their own mesenchymes, the Wolffian duct mesenchyme (WDM) and Müllerian duct mesenchyme (MDM). The WDM and MDM are localizated in the mesometrial and anti-mesometrial side of the mesonephros, respectively. On E16.5, the WD epithelium in the XX embry degenerates in the absence of androgen while the WD mesenchyme (WDM) is maintained and differentiates into a part of the mesenchymal compartment of the oviduct and uterus on E18.5 (Birth). During the female fate differentation from E14.5 and E16.5 in the XX embryo, the WDM undergoes cellular remodeling and increases chromatin accessibility in regions enriched with estrogen response element (ERE+). The WDM also produces IGF1 transiently to facilitate the growth of the MD. In the E18.5 uterus, the WDM derived cells reside predominatly in the mesometrial side with specific expression of AKAP12 and higher TGFB1 sginaling; on the other hand, the MDM-derived cells are exclusively localized in the anti-mesometrial side with specific expression of HAND2 and higher WNT signaling. In the XY embryo, the WDM differentiates into smooth muscle (SM) cells from E14.5 to E16.5 under the action of androgens. During the male fate differentiation, the WDM in the XY embryo increases chromatin accessibility in regions enriched with androgen response element (ARE+) and express higher smooth muscle differentiation related transcriptional factors (TFs).

### The Wolffian duct mesenchyme gives rise to a distinct mesenchymal population for uterine formation after sexual differentiation of reproductive tracts

Despite of the critical roles of mesenchyme in female reproductive tract development, our understanding about the embryonic origin of its mesenchymal tissues has not been complete.

When using *Amhr2-cre; Rosa-LacZ* mouse line to track the Müllerian duct mesenchyme that expresses *Amhr2* in the female mouse embryo, all the LacZ positive mesenchymal cells are located at the anti-mesometrial side of the uterus and never observed in the mesometrial side (*61*). Our results from the *Amhr2-Cre; Rosa-Tomato* model also confirm this observation. This absence of *Amhr2^+^*cells in the mesometrial side cannot be due to inefficient Cre activities because the *Cre* is under the control of *Amhr2* promoter and endogenous *Amhr2* expression is also absent at the mesometrial side (*22*). These results demonstrate that the entire *Amhr2^+^* Müllerian duct mesenchyme contributes to mesenchymal tissues solely at the anti-mesometrial side of the uterus. As a result, when using *Amhr2-Cre* to ablate critical genes for female reproductive tract development and function, the ablation and phenotype only occurs in the anti-mesometrial side (*62–64*). For example, when β-catenin, the intracellular signaling transducer of the canonical WNT pathway, was inactivated in the *Amhr2-Cre; Ctnnb1-flox* conditional knockout mouse, only at the anti-mesometrial side did mesenchymal cells become deficient in smooth muscle differentiation. Those at the mesometrial side remained intact and differentiated normally into myometrium (*62, 63*). These observations imply the presence of a distinct mesenchymal populations in the mesometrial side of the uterus.

Using in vivo genetic lineage tracing, we reveal that the Wolffian duct mesenchyme gives rise to these mesenchymal cells at the mesometrial side of the uterus. Transcriptomic analysis and specific gene expression examination reveal gene expression similarities and differences between mesenchymal populations at the mesometrial and anti-mesometrial side of the uterus at birth. Both mesenchymal populations express typical mesenchymal markers (*Vim*, *Col1a1*, *Col1a2* and *Col3a1*) and show enrichment of angiotensin and estrogen signaling in higher expressed genes in them. However, compared to the *Amhr2^+^*Müllerian duct mesenchyme, the *Gli1^+^* Wolffian duct mesenchyme seems to be conditioned with smooth muscle differentiation signaling. For example, *Tgfβ1* signaling, one major driver for smooth muscle differentiation (*65*), and its downstream gene AKAP12 is enriched only in the *Gli1^+^* Wolffian duct mesenchyme. On the other hand, the Müllerian duct mesenchyme specific gene *Hand2* is exclusively expressed in the stroma and remains absent in the smooth muscle layer in the postnatal uterus (*66*).

We also found that, after birth, the localization of Wolffian duct mesenchyme-derived cells remain at the mesometrial side of the uterus. It is well-noted that the mesometrial and anti-mesometrial mesenchyme in the uterus exhibit differential expression in postnatal development and function. In postnatal development, the *Amhr2^+^* Müllerian duct mesenchyme (giving rise to the anti-mesometrial mesenchyme) had higher WNT signaling activity, which is essential for limiting gland formation to the anti-mesometrial side of the uterus (*49*). During pregnancy, estrogen and progesterone receptors coordinate actions of estrogen and progesterone in the preparation of the uterus for implantation and placentation, which occur at the anti-mesometrial and mesometrial side, respectively (*67, 68*). Interestingly, dynamic and differential expression of estrogen receptor α (ESR1) and progesterone receptor (PGR) are observed between mesenchymal cells at the mesometrial and anti-mesometrial poles during pregnancy (*69, 70*). Multiple single cell RNA transcriptomic studies have revealed mesenchymal heterogeneity in the uterus and have identified several stromal populations in the uteri at the age of postnatal 6 (2 mesenchymal populations, inner and outer stromal populations) (*71*), postnatal 12 (4 mesenchymal populations) (*72*) and in the adult uterus on the estrus stage (3 populations of fibroblasts and 2 population of perivascular cells) (*73*). However, the spatial expression of marker genes for these mesenchymal populations along the mesometrial-anti-mesometrial axis were not examined in these studies. Our discovery of the differential embryonic origins of mesometrial and anti-mesometrial mesenchyme in the uterus has potential implications in the differential gene expression along the mesometrial-anti-mesometrial pole for gland formation and uterine functions.

### The transcriptome and chromatin accessibility analyses during sexual differentiation highlight how hormonal environment alters the fate of the Wolffian duct mesenchyme

The contribution of Wolffian duct mesenchyme to the female reproductive tract demonstrates that the Wolffian duct mesenchyme is bipotential with sexually dimorphic differentiation capacity. Androgen is known to be the predominant hormone for masculinizing Wolffian ducts and promoting the differentiation of the Wolffian duct mesenchyme into smooth muscle cells in the male embryo (*18, 34*). Our observations of myogenesis as the top signaling pathway in the upregulated genes in the Wolffian duct-dervied mesenchymal cells and increased chromatin accessibilities in genomic regions harboring the top motif ARE corroborate with this notion. On the other hand, in the female embryo where androgens are absent, the differentiation of the Wolffian duct mesenchyme into smooth muscle cells does not occur. Although these mesenchymal cells remain undifferentiated in the female embryo, their chromatin accessibility increases in genomic regions containing multiple transcriptional factor motifs with estrogen receptor α (ESR1) as the top one. These observations suggest the estrogen action may play a role in modulating epigenetic landscape in the Wolffian duct mesenchyme in the female embryo. The existence of the ESR1 action in the fetal reproductive tract tissues in the female embryos has been supported by the expression of *Esr1* (*74*) and ESR1 (*75*) in the mesonephric mesenchyme and detection of widespread ER transcriptional activities in the ERE-luciferase reporter mouse (*76*). Consistent with these results on the ESR1 action, we observed increased expression of *Igf1,* a classic ESR1 transcriptional target gene in the mesometrial mesenchyme of the uterus (*77*). Despite the action of ESR1 in the female embryo, the *Esr1* knockout female mouse appears to have normal morphology and histology of the female reproductive tract (*44*). However, a recent study using the aromatase knockout mouse (the critical enzyme for estrogen production) demonstrates that the lack of peri- and postpubertal estrogen diminished uterine responses to estrogen later in life (*78*). Therefore, these observations raise the possibility that the fetal estrogen/ESR1 signaling plays a role in establishing epigenetic landscape for optimizing the Wolffian duct mesenchyme’s responsiveness to estrogen when it is incorporated into the female reproductive tract.

### The Wolffian duct mesenchyme provides critical paracrine growth factors for Müllerian duct development after sexual differentiation of reproductive tracts

Extensive studies have focused on the functional significances of the Wolffian duct in early formation of the Müllerian duct. However, our results indicate that in the female embryo, the remaining mesenchyme of the Wolffian duct still plays a role after Wolffian duct regression and provides critical paracrine growth factors for the Müllerian duct growth. Although we focused on *Igf1 that* is expressed higher in mesometrial side where the *Gli1^+^* Wolffian duct mesenchyme are localized, IGF1 supplementation did not completely rescue the phenotype of decreased lumen expansion in the ablation group. These results indicate other paracrine growth factors from the Wolffian duct mesenchyme may also contribute to the fetal growth of the Müllerian duct. A complementary experiment that ablates the *Amhr2^+^* Müllerian duct mesenchyme in the female reproductive tract was performed by ectopic postnatal AMH administration (PND1-6) to the female rats (*71*). The AMH treatment inhibited *Amhr2^+^* subluminal mesenchyme expansion and led to stromal hypoplasia and dysregulated mesenchymal paracrine signals. Consequently, these treated female rats failed to develop uterine gland formation and were infertile at the adulthood. It would be intriguing to investigate the impact of the loss of the Wolffian duct mesenchyme on postnatal development and function of the female reproductive tract using our *Gli1^+^*cell ablation model. However, the expression of *Gli1* in other tissues at the fetal stage (*51*) and its ubiquitous expression in the mesenchyme of the female reproductive tract after birth (*79*) preclude the use of our *Gli1-CreER; Rosa-DTA* model from specifically ablating the Wolffian duct mesenchyme or its derived cell to determine its postnatal functions.

To put our discovery in the context of human diseases, the most common gynecological benign tumor in women of reproductive ages is uterine fibroids also known as leiomyomas, which derive from genetically altered mesenchymal cells in the female reproductive tract (*80, 81*). Interestingly, in two uterine fibroid mouse models where gain-of-function mutant forms of *Med12* (*82*) or *Ctnnb1* (*83*) were induced in the *Amhr2^+^* Müllerian duct mesenchyme (giving rise to anti-mesometrial mesenchyme), large leiomyoma nodules grew at the mesometrial side of the uterus. These observations suggest the potential involvement or susceptibility of the Wolffian duct mesenchyme-derived cells or their microenvironments for facilitating the formation/growth of leiomyomas. Therefore, our discovery of the contribution of the Wolffian duct mesenchyme to the female reproductive tract not only provide new perspectives on sexual differentiation of reproductive tracts but also potentially promote our understanding of mesenchyme-associated gynecological complications in the female reproductive tract.

## Methods and Materials

### Animals

*Gli1-CreER* knock-in mice (stock# 007913) on mixed genetic backgrounds (Swiss Webster and C57BL/6J), *Rosa-tdTomato* (stock# 007909) on the C57BL/6J genetic background, *Rosa-DTA* (stock# 006331) on mixed genetic backgrounds (C57BL/6J and CD1), *Gli1-LacZ* (#008211) on mixed genetic backgrounds (Swiss Webster and 129S1/SvImJ) were purchased from the Jackson Laboratory (Bar Harbor, ME). *Amhr2-Cre* were derived from previously-described colony (*84*) and maintained on C57BL/6J genetic background. CD-1 mice were from in-house CD-1 colony. Timed mating was produced by housing two or three females with a male. Vaginal plugs were checked daily and the day when the vaginal plug was found was designated as embryonic day E0.5. All animal procedures were approved by the National Institute of Environmental Health Sciences (NIEHS) and the University of Wisconsin-Madison (UW-Madison) Animal Care and Use Committees and are in compliance with our NIEHS and UW-Madison approved animal study proposals and public laws. Genotyping was determined by Transnetyx or PCR based on genotyping protocols provided by the Jackson Laboratory. All experiments were performed on at least three animals for each genotype.

### Tamoxifen treatment

CreER activity was induced by IP injection of tamoxifen (T-5648, Sigma-Aldrich) per mouse in corn oil, receptively. The dose and timing of tamoxifen treatment were described in the results section. The specificity of *Gli1-CreET; Rosa-Tomato* genetic lineage tracing were confirmed in our previous study (*85*).

### C-section and pup fostering

Tamoxifen injection during pregnancy results in dystocia. To avoid the loss of pups due to dystocia, C-section was performed on the expected day of delivery (E18.5 or E19.5) following NIEHS SOP on Caesarian Section Rederivation (Terminal). Briefly, forceps and blunt scissors were used to incise the uterus, rupture amniotic sacs, and clamp the cord of the pups. Once all pups were free of the uterus, fluid from the nose and mouth of each pup were cleared using sterile cotton swabs. Pups were stimulated to breathe by rolling swabs length wise along the pup from mouth to anus. When all pups were breathing regularly without stimulation, and pup color was pink, the rederived pups were fostered to the CD-1 mom whose pups were removed. To ensure the success of fostering, CD-1 foster moms who delivered less than 3 days ago were used. The fostered pups were monitored closely for any signs of abandonment or cannibalism.

### LacZ staining

The LacZ staining solution was made by dissolving X-gal (Invitrogen) into dimethylformamide to make 40 mg/ml stock solution. The working solution was further prepared by diluting the stock solution to 1 mg/ml in pre-warmed tissue stain base solution (Chemicon). Fresh tissues haboring *Amhr2-Cre; Rosa-Tomato; Gli1-LacZ* were fixed in 4% paraformaldehyde in 1xPBS at 4 °C for 1 h and then stained in the LacZ staining solution at 37 °C for 1–2 h followed by further fixation in 4% paraformaldehyde/PBS at 4 °C overnight.

### Immunofluorescence

Tissues were fixed in 4% paraformaldehyde at 4°C overnight. The tissues and sections were processed for immunostaining as previously described (*52*). Tissues were dehydrated, embedded, and cyrosectioned at 10 μm. The sections were treated for antigen retrieval using commercial antigen unmasking solution (H-3300, VECTOR) and underwent immunostaining procedures. The following primary antibodies were used: rabbit anti-PAX2 (1:200, PRB-276P, Covance), rabbit anti-alpha-smooth muscle (1:200, ab5694, Abcam), mouse anti-vimentin (1:300, Abcam, ab8978), rabbit anti-AKAP12 (*86*) (1:500, a gift from Irwin H. Gelman, Roswell Park Cancer Institute), rabbit anti-Cleaved PARP1(1:300, Abcam, ab32064) . The secondary antibodies conjugated with different fluorescent dyes were used (1:200): Alexa Fluor@ 647 donkey anti-mouse IgG, Alexa Fluor@ 488, 568 or 647 donkey anti-rabbit IgG (Invitrogen). All the sections were imaged under a Leica confocal microscope.

For quantifying the number of cleaved PARP1+ cells (the apoptotic cells), at least 8 serial sections with 75 μm apart between each section from each cultured tissue were pictured.

### Immunohistochemistry

Frozen sections were treated for antigen retrieval using commercial antigen umasking solution (H-3300, VECTOR). Endogenous peroxidase was inactivated with 3% H_2_O_2_ (H325, Fisher Scientific). Sections were incubated with blocking reagent, with primary antibodies goat polyclonal anti-HAND2 (Santa Cruz, sc-9409, a gift from Francesco DeMayo’s lab at NIEHS) at 4 °C overnight. Sections was washed three times, and then incubated with biotinylated anti-rabbit secondary antibody (#94583, Jackson ImmunoResearch) for 30 min at room temperature.

Sections were then incubated with ABComplex/HRP (Vectastain ABC kit, VECTOR), and DAB substrate (SK4100, VECTOR), counterstained with hematoxylin (HHS16, Sigma), and mounted for imaging.

### RNAscope

RNAscope was performed in formalin fixed paraffin embedded tissues using RNAscope® 2.5 HD Detection Reagent (for the detection of *Igf1*) and RNAscope Duplex Detection Kit (for dual detections of *Amhr2* and *Gl1i*) according to the manufacturer’s instructions (Advanced Cell Diagnostics, Newark, CA). Tissues were fixed in fresh 10% formalin for 24h at the room temperature. After fixation, tissues were processed, embedded in paraffin, sectioned at 5 μm, deparaffinized, and rehydrated. Then sections were treated with antigen retrieval buffer and proteinase. Sections were exposed to *Igf1* (443901) or *Amhr2* (489821) + *Gli1* (311001-C2) probes and incubated at 40°C in a hybridization oven for 2 h. Following a series of rinsing, for the single detection of *Igf1*, the signaling is amplified and stained with a red substrate; for the dual detection, *Amhr2* and Gli1 signaling were amplified using amplifier conjugated horseradish peroxidase and alkaline phosphatase, respectively.

Then, sections were incubated sequentially with a green and then a red substrate solution to generate chromogenic colors for *Amhr2* and *Gli1* expression, respectively. After the staining, all the sections were counter-stained with Gill’s hematoxylin I, air-dried and mounted. All sections are imaged under a Zeiss compound microscope.

### Ex vivo organ culture

E14.5 mesenephroi with ovaries attached were cultured at 37°C with 5% CO_2_/95% air on MilliCELL-CM culture plate insert 0.4 μm filters (Millipore) in Dulbecco’s Minimal Eagle Medium: Nutrient Mixture F-12 (DMEM/F12) supplemented with 10% fetal calf serum (Hyclone), and 100 U/ml Penicillin-Streptomycin in the presence of and 500 nM 4-hydroxytamoxifen (H7904, Sigma) and 200ng/ml SHH (464-SH-025, R&D system). The hydroxytamoxifen is the active metabolite of tamoxifen in vivo for inducing the nuclear translocation of CreER to elicit gene recombination (*87*). The supplementation of SHH enhanced CreER expression and thus the CreER mediated ablation efficiency. After 48h of culture in the presence of SHH and 4-hydroxytamoxifen, tissues were cultured for another 48h with or without IGF1 (291-G1-200, R&D system). After culture, the whole tissues were imaged under a fluorescence stereo microscope for *tdTomato* expression and bright-light images. Image J was used to quantify the corrected total fluorescence (*88*) and lumen areas of the whole mount tissues.

### Fluorescence-activated cell sorting

After tamoxifen treatment at the dose of 1 mg per dam on E12.5 and E13.5, tomato+ cells were isolated from E14.5 and E16.5 *Gli1-CreER+; Rosa-tdTomato+* mesonephroi, E19.5 *Gli1-CreER+;Rosa-tdTomato+* uterus or E19.5 *Amhr2-Cre+;Rosa-tdTomato+* uterus using the following protocol for tissue dissociation. Tissues were dissected in cold 1xPBS and enzymatically dissociated in TrypLE Express (Gibco, 12604-013 -no phenol red) at 37 °C for 20–30 minutes with 1000 rpm shaking in a VorTemp™ 56 Shaking Incubator (Labnet). Cells were further dissociated mechanically using a P200 pippette. After quenching enzyme activities by addition 2 volume of sorting buffer (1% FBS in 1xPBS), cell were pelleted for 10 min at 500 g at 4 °C and suspended in sorting buffer, which was added through the cell strainer in the cap (remove tissue clogs) to 5 ml polystyrene round-bottom tube (#352235, Falcon).

Cell sorting was performed on a BD FACS Aria II in the NIEHS Flow Cytometry Center and tdTomato+ cells were collected in 5 ml polystyrene round bottom tube that had been coated in sorting buffer overnight.

### RNA extraction and RNA-seq

Each group in RNA-seq experiments had three biological replicates with each pooled from cells in two or three times cell sortings. RNA extractions were performed using PicoPure RNA Isolation kit (Life technologies, USA) according to the manufacturer’s protocol. The quality of RNA was measured in a 2100 Bioanalyzer Instrument (Agilent) with bioanalyzer high-sensitivity RNA kits according to the manufacturer’s protocol. The RNA integrity number (RIN) of RNAs were all above 9.7. The RNA concentration was measured in a Qubit Fluorometer (ThermoFisher Scientific). For sorted Gli1^+^ mesenchymal cells, 250 ng RNA were used to generate libraries using Truseq RNA non-stranded kit (Illumina), which were sequenced in NextSeq 500 platform at NIEHS Epigenomics Core with the sequencing parameter, single end 75 nt reads. For sorted *Gli1^+^* and *Amhr2^+^* cell from E19.5 uterus, TruSeq Stranded mRNA kit was used to generate libraries that were sequenced by ActiveMotif on the Illumina platform with the sequencing parameter, paired end 75 nt reads.

### ATAC-seq

ATAC-seq was performed with Illumina Nextera DNA preparation kit (Illumina, FC-121-1030) according to the previous protocol (*89*). Two replicates were included in each group. 20,000 sorted tdTomato+ cells from *Gli1-CreER+;Rosa-tdTomato+* E14.5 and E16.5 mesonephroi were collected, and permeabilized with ice cold lysis buffer. The transposition reactions were carried out at 37 °C for 30 min in 50 μl volume containing (25 μl 2x TD buffer, 2.5 μl transposase (100 nM final), 16.5 μl PBS, 0.5 μl 1% digitonin, 0.5 μl 10% Tween-20, 5 μl H_2_O). Digested DNA was purified with Zymo DNA Clean and Concentrator-5 Kit (cat# D4014). Library pre-amplification was done with KAPA HiFi HotStart ReadyMixPCR Kit. Thermocycler conditions were 72°C for 5 min, 98°C for 30 sec, followed by 5 cycles of [98°C for 10 sec, 63°C for 30 sec, 72°C for 1 min] then hold at 4°C. The qPCR amplification using 5 ul of pre-amplified mixture was performed to determine additional cycles, which were between 5-6. Final amplification was performed with the remainder of the pre-amplified DNA and final PCR products were purified with Zymo DNA Clean and Concentrator-5 Kit and eluted in 20 μl H2O. KAPA Pure beads were used to size select 150-450 bp DNA fragments (x0.6 and x1.5 cut). The size selected library was measured in a Cubic II for concentration and in a Bioanalyzer for size distribution, and sequenced in NextSeq 500 platform at NIEHS Epigenomics Core. Sequencing parameter was paired end 2x35 nt reads.

### Bioinformatic analyses of next generation sequencing data

For analysis of the RNA-seq datasets, raw sequences were filtered to remove all reads with mean base quality score < 20; for paired-end data, both reads were required to pass this filter. Filtered reads were mapped against the mm10 reference assembly by STAR v2.5 (*90*) with parameters "--outSAMattrIHstart 0 --outFilterType BySJout --alignSJoverhangMin 8 -- limitBAMsortRAM 55000000000 --outSAMstrandField intronMotif --outFilterIntronMotifs RemoveNoncanonical". Counts per gene were determined via featureCounts (Subread v1.5.0-p1) (*91*) with parameters "-s0" for the unstranded single-end data or "-s2 -Sfr -p" for the stranded paired-end data. Evaluated gene models are GENCODE VM18 annotations, as defined in the wgEncodeGencodeBasicVM18 table downloaded from the UCSC Table Browser (https://genome.ucsc.edu/cgi-bin/hgTables) on December 17, 2018. Entries were associated with Entrez gene identifiers when possible. Differential expression analysis was performed with DESeq2 v1.14.1 (*92*) in R v3.3.2. Differentially expressed gene thresholds were set at FDR 0.05, fold change 1.5, and minimum average TPM 1. Pathway analysis was performed in Enrichr (*93*) by job submission at https://maayanlab.cloud/Enrichr/. Presented results are from MSigDB Hallmark 2020, with significant pathways identified at adjusted p-value < 0.05. For the E19.5 Gli1+ and Amhr2+ mesenchyme dataset, upstream regulatory analysis was performed according to the (Ingenuity Pathway Analysis) IPA guidelines (*94*).

For analysis of the ATAC-seq dataset, raw sequences were filtered to remove all pairs with mean base quality score < 20 for either read. Filtered read pairs were mapped against the mm10 reference assembly by Bowtie v1.2 (*95*) with parameters "-m 1 -X 2000 --chunkmbs 1024". Hits to chrM were discarded via samtools v1.3.1 (*96*). Duplicate mapped read pairs were removed by Picard tools MarkDuplicates.jar (v1.110) (http://broadinstitute.github.io/picard). Downstream analysis considered only the 9bp at the 5’ end of each read. Peak calls per sample were made by MACS2 v2.1.1 (*97*) with parameters "callpeak -g mm -q 0.0001 --keep-dup=all -- nomodel --extsize 9", followed by combining nearby peaks via BEDTools v2.24.0 (*98*) "merge -d 200". From the initial peak sets, a single set of unified peaks was generated to facilitate comparisons across samples. The unified peaks were defined by collapsing peaks from all samples and collecting regions that overlapped a called peak from 2 or more samples. Nearby unified peaks were combined via BEDTools v2.24.0 "merge -d 200", then filtered to require a minimum length of 50bp. ATAC-seq signal per peak was determined with BEDTools v2.24.0 coverage with the "-counts" option. EdgeR v3.16.5 (*99*) in R v3.3.2 was used to identify differential ATAC-seq signal between sample groups at FDR 0.01. Differential peaks were stratified into TSS proximal or TSS distal subsets based on a distance cutoff of 1Kb.Enriched motif analysis was performed by HOMER v4.10.3 (*100*) findMotifsGenome.pl with "-size given" after expanding differential peaks to a minimum width of 200bp; results presented here are for queries limited to peaks more than 1Kb from the nearest annotated TSS (distance measured after expansion to minimum 200bp size).

PCA plots were generated with the plotPCA function in DESeq2 v1.14.1, using the default setting of 500 most variant entries (either genes or ATAC-seq peak regions).

### Statistical analyses

A minimum of three biological replicates were used in each examined point in lineage tracing, immunofluorescence, immunohistochemistry, and RNA-scope. Quantitative data is presented as mean±SEM and the sample sizes are indicated in figure legends. One way ANOVA test was used for evaluating significant differences among three groups in Figure 4C&4D. Two-tail Student’s t test was used for evaluating significant differences between control and ablation groups in supplementary figure 5E. The significance level was set at p<0.05.

## Acknowledgments

We are thankful to the NIEHS Epigenomics and DNA Sequencing Core for the RNA-seq and ATAC-seq sequencing; Maria Sifre at the NIEHS Flow Cytometry Center for her help with cell sorting; Comparative Medicine Branch for mouse colony maintenance; Dr. Hongyao Yu in Dr. Guang Hu’s Group at the NIEHS for his assistance with ATAC-seq; and Paula Brown and Karina Rodriguez in Dr. Yao’s lab for their help with IPA analysis and preparing and shipping samples and mice.

## Funding Sources

Intramural Research Program of National Institute of Environmental Health Sciences Z01-ES102965 (HHCY)

National Institute of Child Health and Development R00-HD096051 (FZ)

## Author Contributions

FZ performed most of the experiments; SJ performed staining and quantifications of lumen areas and fluorescent signaling in Figure 4 and Supplementary figure 5; FZ and HHCY designed the study, analyzed the data, and wrote the paper; SAG analyzed the next-generation sequencing data and edited the paper.

## Data and materials availability

All data are available in the main text or the supplementary materials upon reasonable request. RNA-seq and ATAC-seq data have been deposited in the GEO database under the accession code GSE179876.

