## Supplementary figures and tables for "Contribution of the Wolffian duct mesenchyme to the formation of the female reproductive tract"

**Fig. S1.**

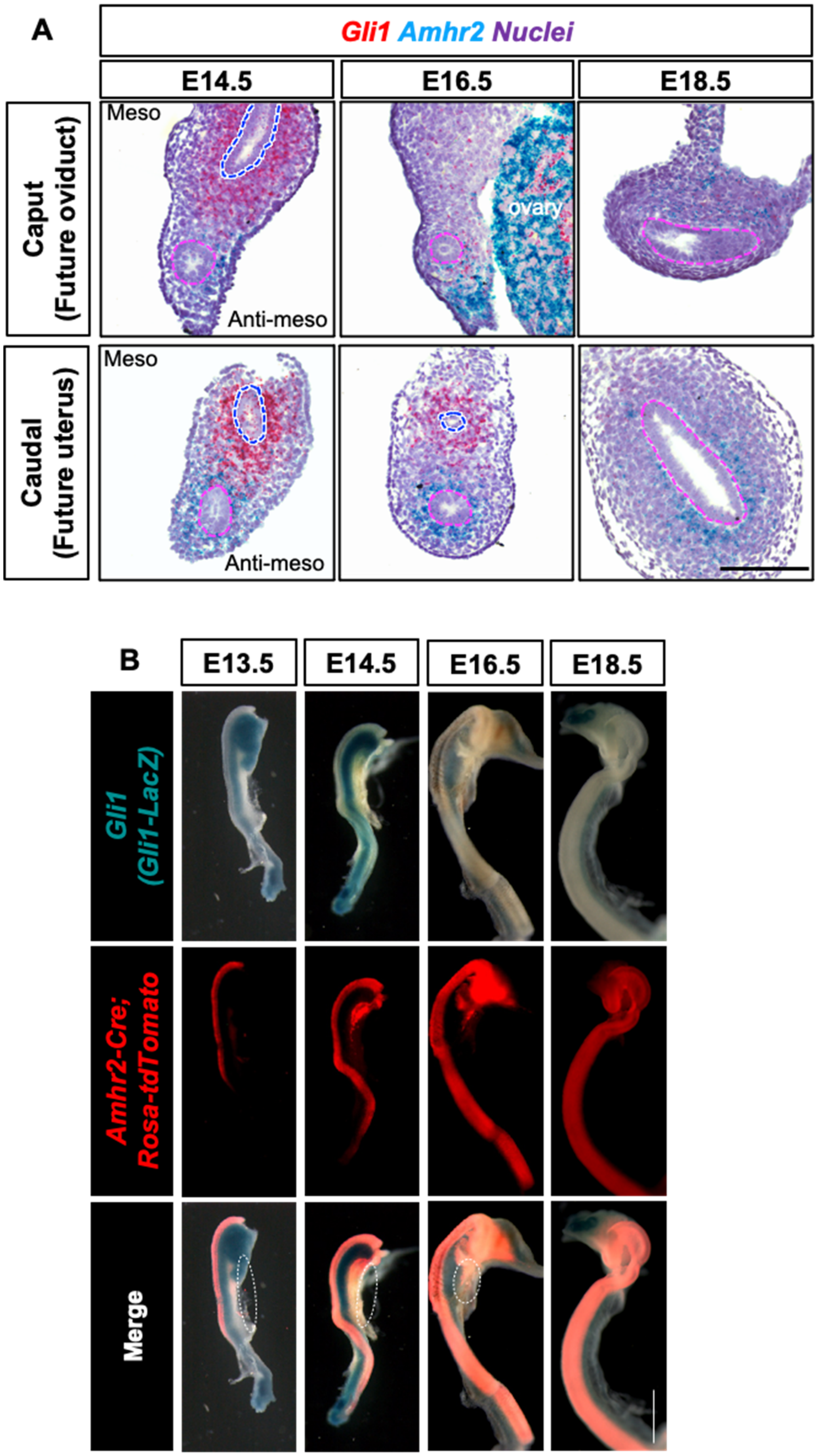
A
Gli1 Amhr2 Nuclei
E14.5
E16.5
E18.5
Caput (Future oviduct)
Meso
Anti-meso
ovary
Caudal (Future uterus)
Meso
Anti-meso
B
E13.5
E14.5
E16.5
E18.5
Gli1 (Gli1-LacZ)
Amhr2-Cre; Rosa-tdTomato
Merge

(**A**) Duplex RNAscope assay to co-detect *Amhr2* (Green) and *Gli1*(Red) in fetal reproductive tracts in XX embryos on E14.5, E16.5 and E18.5. Nuclei is stained with hematoxylin (Blue). All images are oriented with the mesometrial side of the tissues up and the anti-mesometrial side down. Pink and blue dashed lines circle Müllerian and Wolffian ducts, respectively. Scale bar: 100 µm. (**B**) *Amhr2-Cre; Rosa-tdTomato; Gli1-lacZ* double reporter mouse line for the co-detection of *Gli1* (Blue) and *Amhr2 lineage* (Red). Scale bar: 0.5 mm. White dashed lines indicates the ovairies. N=3 in each time point examined.

**Fig. S2.**

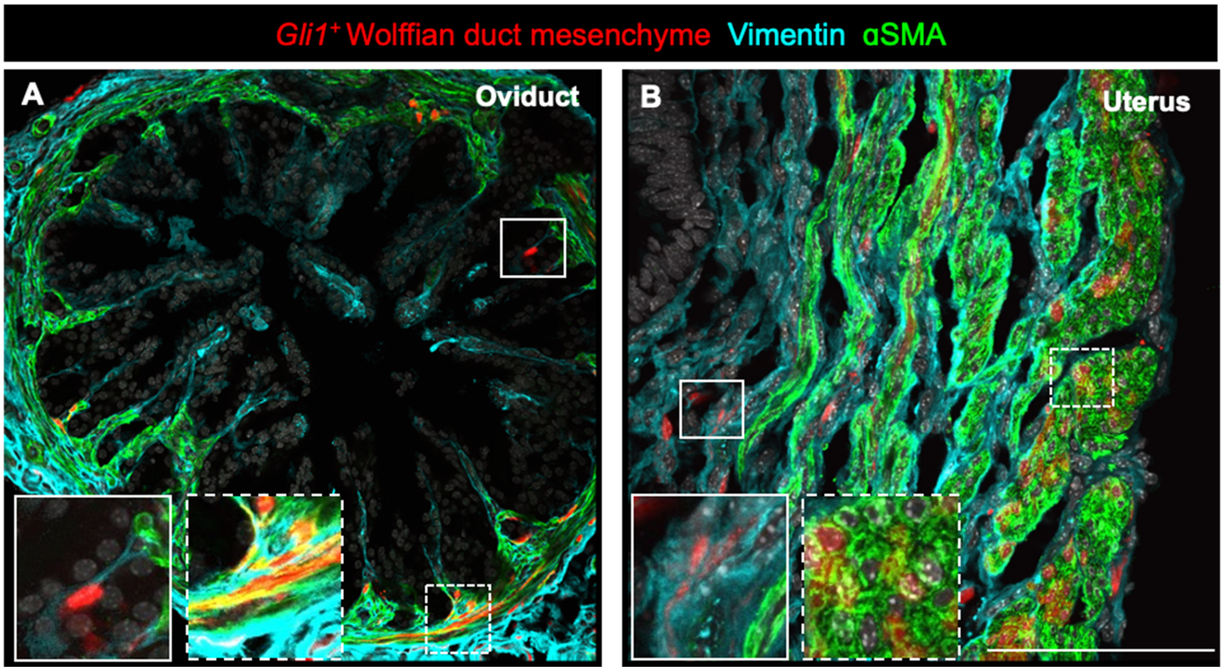


**(A & B)** Immunofluorescence section staining of fibroblast and smooth muscle markers (vimentin and αSMA, respectively) in the adult oviduct (A) and adult uterus (B) from the *Gli1-CreER; Rosa-tdTomato* lineage tracing model in the Figure 1. Boxes outlined with solid and dashed lines indicate the co-expression of tdTomato with vimentin and αSMA, respectively, in higher magnifications. Scale bar: 100 µm. N=5.

**Fig. S3.**

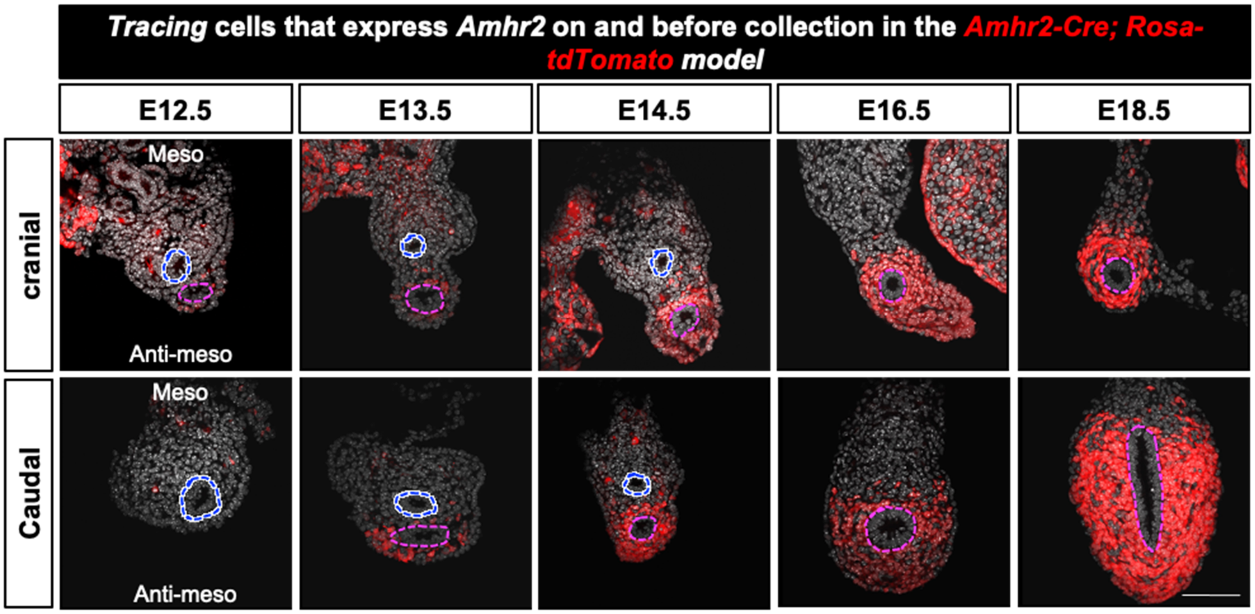


Detection of *tdTomato* labeled mesenchymal cells that express *Amhr2* on and before the collection date in cranial and caudal sections of the female reproductive tracts in the *Amhr2-Cre; Rosa-tdTomato* model. All images are oriented with the mesometrial side of the tissues up and the anti-mesometrial side down. Pink and blue dashed lines circle Müllerian and Wolffian ducts, respectively. Scale bar: 50 µm. N=3 in each time point examined.

**Fig. S4.**

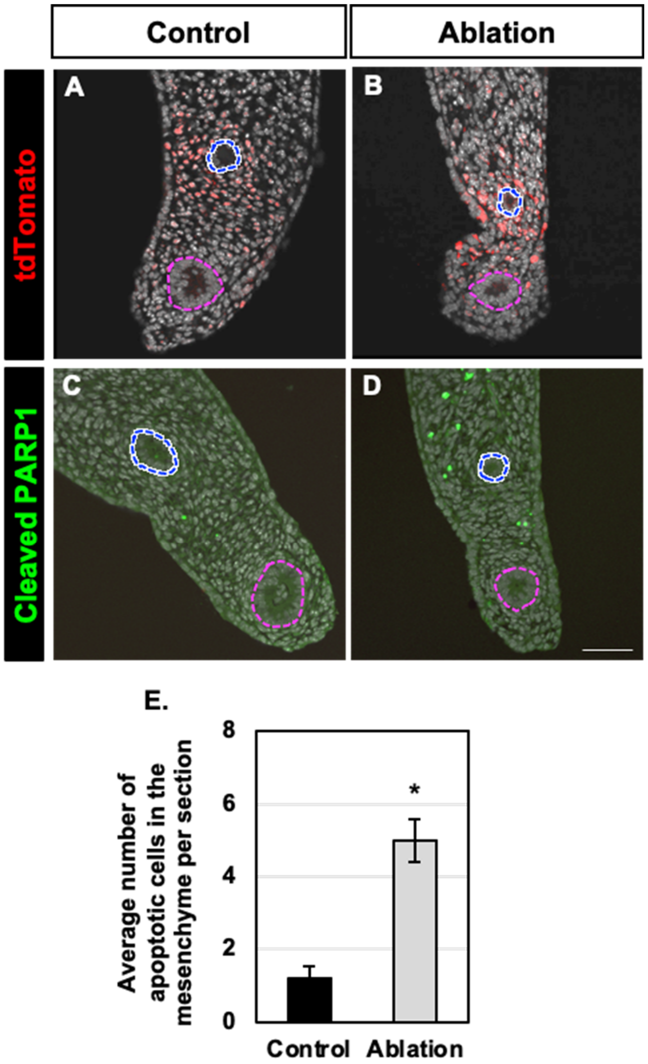


Detection of *tdTomato* labeled *Gli1*$^{+}$ Wolffian duct mesenchyme and apoptotic cells in fetal female reproductive tract after 2-day culture in the control and ablation groups. (**A & B**) Detection of tdTomato in the control and ablation groups. (**C & D**) Detection of the apoptotic marker, cleaved PARP1 in the control and ablation groups. (**E**) Average number of apoptotic cells in the mesenchyme per section. N=5 in both control and ablation groups. *: P<0.05. All images are oriented with the mesometrial side of the tissues up and the anti-mesometrial side down. Pink and blue dashed lines circle Müllerian and Wolffian ducts, respectively. Scale bar: 50 µm.

**Table S1.**

| *type* | *comparison* | *total differential peaks* | *TSS proximal* | | *TSS distal* | |
|---|---|---|---|---|---|---|
| E14.5, compare sexes | E14F > E14M | 1,476 | 67 | 4.5% | 1,409 | 95.5% |
| | E14M > E14F | 2,523 | 99 | 3.9% | 2,424 | 96.1% |
| E16.5, compare sexes | E16F > E16M | 3,207 | 460 | 14.3% | 2,747 | 85.7% |
| | E16M > E16F | 9,891 | 498 | 5.0% | 9,393 | 95.0% |
| female, compare ages | E16F > E14F | 5,935 | 2,360 | 39.8% | 3,575 | 60.2% |
| | E14F > E16F | 13,912 | 939 | 6.7% | 12,973 | 93.3% |
| male, compare ages | E16M > E14M | 8,158 | 700 | 8.6% | 7,458 | 91.4% |
| | E14M > E16M | 10,292 | 812 | 7.9% | 9,480 | 92.1% |

The number of ATAC-seq peaks with differential signal in pairwise group comparisons. Differential signal threshold is FDR 0.01 in EdgeR. TSS proximal is defined as within 1Kb of TSS.

**Table S2.**

| Rank | Name | log P-pvalue | % of Targets Sequences with Motif | % of Background Sequences with Motif |
|---|---|---|---|---|
| 1 | GRE(NR),IR3/RAW264.7-GRE-ChIP-Seq(Unpublished)/Homer | -2.33E+03 | 52.04% | 4.37% |
| 2 | GRE(NR),IR3/A549-GR-ChIP-Seq(GSE32465)/Homer | -2.22E+03 | 42.10% | 2.35% |
| 3 | ARE(NR)/LNCAP-AR-ChIP-Seq(GSE27824)/Homer | -1.96E+03 | 51.63% | 5.78% |
| 4 | PGR(NR)/EndoStromal-PGR-ChIP-Seq(GSE69539)/Homer | -1.91E+03 | 47.83% | 4.78% |
| 5 | PR(NR)/T47D-PR-ChIP-Seq(GSE31130)/Homer | -7.61E+02 | 81.96% | 43.56% |
| 6 | NF1(CTF)/LNCAP-NF1-ChIP-Seq(Unpublished)/Homer | -3.14E+02 | 19.44% | 5.11% |
| 7 | Tcf21(bHLH)/ArterySmoothMuscle-Tcf21-ChIP-Seq(GSE61369)/Homer | -2.46E+02 | 33.72% | 15.57% |
| 8 | TCF4(bHLH)/SHSY5Y-TCF4-ChIP-Seq(GSE96915)/Homer | -1.99E+02 | 47.67% | 28.65% |
| 9 | NeuroG2(bHLH)/Fibroblast-NeuroG2-ChIP-Seq(GSE75910)/Homer | -1.97E+02 | 47.30% | 28.38% |
| 10 | ZBTB18(Zf)/HEK293-ZBTB18.GFP-ChIP-Seq(GSE58341)/Homer | -1.94E+02 | 22.04% | 8.89% |
| 11 | Atoh1(bHLH)/Cerebellum-Atoh1-ChIP-Seq(GSE22111)/Homer | -1.85E+02 | 35.41% | 18.93% |
| 12 | AR-halfsite(NR)/LNCaP-AR-ChIP-Seq(GSE27824)/Homer | -1.75E+02 | 81.26% | 64.12% |
| 13 | Ap4(bHLH)/AML-Tfap4-ChIP-Seq(GSE45738)/Homer | -1.55E+02 | 35.66% | 20.38% |
| 14 | Olig2(bHLH)/Neuron-Olig2-ChIP-Seq(GSE30882)/Homer | -1.50E+02 | 54.07% | 36.95% |
| 15 | Tcf12(bHLH)/GM12878-Tcf12-ChIP-Seq(GSE32465)/Homer | -1.47E+02 | 27.86% | 14.52% |
| 16 | NeuroD1(bHLH)/Islet-NeuroD1-ChIP-Seq(GSE30298)/Homer | -1.36E+02 | 26.95% | 14.26% |
| 17 | Myf5(bHLH)/GM-Myf5-ChIP-Seq(GSE24852)/Homer | -1.29E+02 | 22.66% | 11.30% |
| 18 | Pitx1:Ebox(Homeobox,bHLH)/Hindlimb-Pitx1-ChIP-Seq(GSE41591)/Homer | -1.25E+02 | 11.02% | 3.68% |
| 19 | NF1-halfsite(CTF)/LNCaP-NF1-ChIP-Seq(Unpublished)/Homer | -1.17E+02 | 41.35% | 27.12% |
| 20 | Ascl1(bHLH)/NeuralTubes-Ascl1-ChIP-Seq(GSE55840)/Homer | -1.12E+02 | 38.34% | 24.77% |

Top known motifs in regions with increased chromatin accessibility (E14.5XY>14.5XX distal peaks)

**Table S3.**

| Rank | Name | log P-pvalue | % of Targets Sequences with Motif | % of Background Sequences with Motif |
|---|---|---|---|---|
| 1 | GRE(NR),IR3/RAW264.7-GRE-ChIP-Seq(Unpublished)/Homer | -3.887e+03 | 30.54% | 3.76% |
| 2 | GRE(NR),IR3/A549-GR-ChIP-Seq(GSE32465)/Homer | -3.446e+03 | 22.81% | 2.04% |
| 3 | PGR(NR)/EndoStromal-PGR-ChIP-Seq(GSE69539)/Homer | -3.192e+03 | 27.77% | 3.89% |
| 4 | ARE(NR)/LNCAP-AR-ChIP-Seq(GSE27824)/Homer | -3.041e+03 | 30.81% | 5.33% |
| 5 | PR(NR)/T47D-PR-ChIP-Seq(GSE31130)/Homer | -1.614e+03 | 69.15% | 40.16% |
| 6 | NF1(CTF)/LNCAP-NF1-ChIP-Seq(Unpublished)/Homer | -1.379e+03 | 21.17% | 5.37% |
| 7 | Tcf21(bHLH)/ArterySmoothMuscle-Tcf21-ChIP-Seq(GSE61369)/Homer | -1.088e+03 | 35.77% | 16.00% |
| 8 | Atoh1(bHLH)/Cerebellum-Atoh1-ChIP-Seq(GSE22111)/Homer | -8.122e+02 | 36.34% | 18.68% |
| 9 | NeuroG2(bHLH)/Fibroblast-NeuroG2-ChIP-Seq(GSE75910)/Homer | -7.391e+02 | 45.85% | 27.30% |
| 10 | TCF4(bHLH)/SHSY5Y-TCF4-ChIP-Seq(GSE96915)/Homer | -7.144e+02 | 45.64% | 27.40% |
| 11 | Tcf12(bHLH)/GM12878-Tcf12-ChIP-Seq(GSE32465)/Homer | -7.128e+02 | 30.98% | 15.45% |
| 12 | Ap4(bHLH)/AML-Tfap4-ChIP-Seq(GSE45738)/Homer | -6.753e+02 | 38.12% | 21.47% |
| 13 | ZBTB18(Zf)/HEK293-ZBTB18.GFP-ChIP-Seq(GSE58341)/Homer | -6.522e+02 | 21.12% | 8.89% |
| 14 | Myf5(bHLH)/GM-Myf5-ChIP-Seq(GSE24852)/Homer | -6.066e+02 | 24.69% | 11.73% |
| 15 | MyoD(bHLH)/Myotube-MyoD-ChIP-Seq(GSE21614)/Homer | -5.776e+02 | 25.40% | 12.50% |
| 16 | Olig2(bHLH)/Neuron-Olig2-ChIP-Seq(GSE30882)/Homer | -5.680e+02 | 51.35% | 34.43% |
| 17 | NeuroD1(bHLH)/Islet-NeuroD1-ChIP-Seq(GSE30298)/Homer | -5.541e+02 | 27.54% | 14.33% |
| 18 | MyoG(bHLH)/C2C12-MyoG-ChIP-Seq(GSE36024)/Homer | -5.107e+02 | 31.04% | 17.54% |
| 19 | Ascl1(bHLH)/NeuralTubes-Ascl1-ChIP-Seq(GSE55840)/Homer | -4.931e+02 | 39.72% | 25.02% |
| 20 | AR-halfsite(NR)/LNCaP-AR-ChIP-Seq(GSE27824)/Homer | -4.615e+02 | 76.39% | 61.73% |

Top known motifs in regions with increased chromatin accessibility (E16.5XY>E16.5XX)

**Table S4.**

| Rank | Name | log P-pvalue | % of Targets Sequences with Motif | % of Background Sequences with Motif |
|---|---|---|---|---|
| 1 | Hoxc9(Homeobox)/Ainv15-Hoxc9-ChIP-Seq(GSE21812)/Homer | -2.18E+02 | 29.38% | 9.69% |
| 2 | LEF1(HMG)/H1-LEF1-ChIP-Seq(GSE64758)/Homer | -1.70E+02 | 33.07% | 13.88% |
| 3 | Tcf3(HMG)/mES-Tcf3-ChIP-Seq(GSE11724)/Homer | -1.53E+02 | 18.31% | 5.32% |
| 4 | CDX4(Homeobox)/ZebrafishEmbryos-Cdx4.Myc-ChIP-Seq(GSE48254)/Homer | -1.47E+02 | 36.41% | 17.45% |
| 5 | NF1(CTF)/LNCAP-NF1-ChIP-Seq(Unpublished)/Homer | -1.39E+02 | 17.25% | 5.13% |
| 6 | Tcf7(HMG)/GM12878-TCF7-ChIP-Seq(Encode)/Homer | -1.36E+02 | 19.73% | 6.60% |
| 7 | PBX2(Homeobox)/K562-PBX2-ChIP-Seq(Encode)/Homer | -1.25E+02 | 32.79% | 15.90% |
| 8 | Pitx1:Ebox(Homeobox,bHLH)/Hindlimb-Pitx1-ChIP-Seq(GSE41591)/Homer | -1.14E+02 | 13.20% | 3.66% |
| 9 | TCF4(bHLH)/SHSY5Y-TCF4-ChIP-Seq(GSE96915)/Homer | -1.10E+02 | 46.49% | 28.11% |
| 10 | Tcf21(bHLH)/ArterySmoothMuscle-Tcf21-ChIP-Seq(GSE61369)/Homer | -1.10E+02 | 31.23% | 15.60% |
| 11 | Atoh1(bHLH)/Cerebellum-Atoh1-ChIP-Seq(GSE22111)/Homer | -1.06E+02 | 35.20% | 18.97% |
| 12 | GATA(Zf),IR3/iTreg-Gata3-ChIP-Seq(GSE20898)/Homer | -1.05E+02 | 11.14% | 2.87% |
| 13 | Fli1(ETS)/CD8-FLI-ChIP-Seq(GSE20898)/Homer | -1.04E+02 | 32.93% | 17.28% |
| 14 | TCFL2(HMG)/K562-TCF7L2-ChIP-Seq(GSE29196)/Homer | -9.95E+01 | 7.81% | 1.49% |
| 15 | GATA(Zf),IR4/iTreg-Gata3-ChIP-Seq(GSE20898)/Homer | -9.80E+01 | 7.67% | 1.45% |
| 16 | NeuroG2(bHLH)/Fibroblast-NeuroG2-ChIP-Seq(GSE75910)/Homer | -9.74E+01 | 45.42% | 28.20% |
| 17 | NeuroD1(bHLH)/Islet-NeuroD1-ChIP-Seq(GSE30298)/Homer | -9.30E+01 | 28.25% | 14.40% |
| 18 | Etv2(ETS)/ES-ER71-ChIP-Seq(GSE59402)/Homer(0.967) | -8.83E+01 | 30.02% | 16.07% |
| 19 | EWS:ERG-fusion(ETS)/CADO_ES1-EWS:ERG-ChIP-Seq(SRA014231)/Homer | -8.21E+01 | 25.90% | 13.32% |
| 20 | ERG(ETS)/VCaP-ERG-ChIP-Seq(GSE14097)/Homer | -8.14E+01 | 42.87% | 27.33% |

Top known motifs in regions with increased chromatin accessibility (E14.5 XX>E14.5 XY)

**Table S5.**

| Rank | Name | log P-pvalue | % of Targets Sequences with Motif | % of Background Sequences with Motif |
|---|---|---|---|---|
| 1 | Pitx1:Ebox(Homeobox,bHLH)/Hindlimb-Pitx1-ChIP-Seq(GSE41591)/Homer | -7.228e+02 | 23.98% | 3.82% |
| 2 | Hoxc9(Homeobox)/Ainv15-Hoxc9-ChIP-Seq(GSE21812)/Homer | -4.253e+02 | 30.50% | 10.31% |
| 3 | NF1(CTF)/LNCAP-NF1-ChIP-Seq(Unpublished)/Homer | -3.775e+02 | 20.70% | 5.47% |
| 4 | TCF4(bHLH)/SHSY5Y-TCF4-ChIP-Seq(GSE96915)/Homer | -3.397e+02 | 51.24% | 27.76% |
| 5 | NeuroG2(bHLH)/Fibroblast-NeuroG2-ChIP-Seq(GSE75910)/Homer | -3.042e+02 | 49.85% | 27.70% |
| 6 | Atoh1(bHLH)/Cerebellum-Atoh1-ChIP-Seq(GSE22111)/Homer | -2.917e+02 | 38.48% | 18.80% |
| 7 | PBX2(Homeobox)/K562-PBX2-ChIP-Seq(Encode)/Homer | -2.747e+02 | 34.80% | 16.48% |
| 8 | Tcf21(bHLH)/ArterySmoothMuscle-Tcf21-ChIP-Seq(GSE61369)/Homer | -2.721e+02 | 33.75% | 15.76% |
| 9 | ZBTB18(Zf)/HEK293-ZBTB18.GFP-ChIP-Seq(GSE58341)/Homer | -1.888e+02 | 20.92% | 8.86% |
| 10 | CDX4(Homeobox)/ZebrafishEmbryos-Cdx4.Myc-ChIP-Seq(GSE48254)/Homer | -1.880e+02 | 35.20% | 19.51% |
| 11 | Olig2(bHLH)/Neuron-Olig2-ChIP-Seq(GSE30882)/Homer | -1.815e+02 | 53.79% | 36.08% |
| 12 | NeuroD1(bHLH)/Islet-NeuroD1-ChIP-Seq(GSE30298)/Homer | -1.741e+02 | 27.99% | 14.34% |
| 13 | Tlx?(NR)/NPC-H3K4me1-ChIP-Seq(GSE16256)/Homer | -1.532e+02 | 16.87% | 7.04% |
| 14 | Ap4(bHLH)/AML-Tfap4-ChIP-Seq(GSE45738)/Homer | -1.408e+02 | 34.44% | 20.76% |
| 15 | ERE(NR),IR3/MCF7-ERa-ChIP-Seq(Unpublished)/Homer | -1.405e+02 | 12.10% | 4.33% |
| 16 | RFX(HTH)/K562-RFX3-ChIP-Seq(SRA012198)/Homer | -1.360e+02 | 5.61% | 1.08% |
| 17 | Rfx2(HTH)/LoVo-RFX2-ChIP-Seq(GSE49402)/Homer | -1.312e+02 | 5.94% | 1.26% |
| 18 | X-box(HTH)/NPC-H3K4me1-ChIP-Seq(GSE16256)/Homer | -1.177e+02 | 6.45% | 1.63% |
| 19 | Pdx1(Homeobox)/Islet-Pdx1-ChIP-Seq(SRA008281)/Homer | -1.062e+02 | 30.98% | 19.46% |
| 20 | Ascl1(bHLH)/NeuralTubes-Ascl1-ChIP-Seq(GSE55840)/Homer | -1.040e+02 | 36.73% | 24.54% |

Top known motifs in regions with increased chromatin accessibility (E16.5 XX>E16.5 XY)

**Data S1.**
Differentially expressed genes in our RNA-seq datasets (separate file)

**Data S2.**
Differentially ATAC peaks (separate file)
